## Supplementary Text for "Probing metazoan polyphosphate biology using *Drosophila* reveals novel and conserved polyP functions"

**Supplementary Information**

**Material and methods**

**Polyphosphate extraction:**

Fly samples were lysed in LETS buffer (10 mM Tris-HCl, pH 8.0, 100 mM lithium chloride, 10 mM EDTA, 0.2% SDS); 200µl LETS buffer was added to 20 third instar larvae, and the larvae were crushed using pestle at room temperature. To precipitate the genomic DNA, acid phenol (pH 4.5), equivalent to the volume of the lysis buffer, was added, and the samples were centrifuged at 18000g for 10 minutes at room temperature. The aqueous phase, which is devoid of genomic DNA, was transferred in a fresh vial and mixed with a double volume of chloroform. The mixture was vortexed for three minutes and centrifuged at room temperature at 18000g for five minutes. The aqueous phase, devoid of proteins, was collected and treated with RNase A at 37℃ for four hours, followed by the addition of an equal volume of 1:1 phenol: chloroform mixture. The solution was vortexed for three minutes and centrifuged at room temperature at 18000g for five minutes. The aqueous phase was collected, and chloroform was added to it in a volume double that of the samples. The mixture was vortexed for three minutes and centrifuged at room temperature at 18000g for five minutes. The aqueous phase was collected and mixed with absolute ethanol in the ratio 1:2.5 (vol/vol), followed by overnight incubation at -80℃. The next day, the polyP was precipitated by centrifugation at 18000g at 4℃ for 30 minutes. Ethanol was decanted, and the transparent polyP pellet was air-dried before dissolving in water.

**Cloning of constructs for protein purification:**

The plasmid pTrc-HisB-*ScPpX1* was a kind gift from Adolfo Saiardi (1). N terminal XPRS tagged PPBD (PPXc) was cloned out from pETM41-PPXc (addgene #38329, (2) ) was cloned into parent vector pGEX-6P2-GST using restriction sites BamHI and XhoI to create pGEX-6P2-GST-XPRS-PPBD (pGEX-6P2-GST-PPBD).

**Protein Purification:**

*Sc*Ppx1 *purification*

*E.coli* BL21(DE3) competent cells transformed with pTrc-HisB-*ScPpx1* were incubated as a primary culture in Luria Bertani (LB) broth containing 100 ug/ml ampicillin overnight at 37℃. The culture was diluted into 1 L LB broth and incubated at 37℃ till $OD_{600}$ reached 0.8. Protein expression was induced with 1 mM IPTG for 4-6 hours at 30℃ with shaking at 180 rpm. The cells were pelleted by centrifugation at 3000g for 15 minutes at 4℃. The cell pellet was resuspended in 12 ml ice-cold lysis buffer (25 mM Tris-HCl pH 8.0, 100 mM NaCl). This was followed by lysis of the cells by sonication (15 minutes, output voltage 20, output control 30, duty cycle 30/40) using Branson Sonifier on ice until the lysate became clear. The lysate was centrifuged at 15000g for 20 minutes at 4℃. The supernatant was transferred to Ni-NTA column (1 mL bed volume, pre-equilibrated with lysis buffer) and incubated in an end-over mixer at 4℃ for two hours. The flow through was collected from the column under gravity at room temperature. The column was sequentially washed with ten column volumes each of wash buffer I (25 mM Tris-HCl pH 8.0, 100 mM NaCl, and 50 mM imidazole), and wash buffer II (25 mM Tris-HCl pH 7.4, 100 mM NaCl, and 75 mM imidazole), and eluted in 5 mL elution buffers each containing 25 mM Tris-HCl pH 7.4, 100 mM NaCl, and imidazole (100 mM, 150 mM, 200 mM, and 300 mM respectively). The eluate pooled together was subjected to size exclusion chromatography on a Sephadex-200 sizing column in buffer containing 25 mM Tris-HCl pH 7.4, 100 mM NaCl to obtain isolated fractions of *ScPpx1* protein and remove contaminating proteins. The purified *Sc*Ppx1 protein was divided into single-use aliquots and stored at -80℃.

*GST, GST-PPBD, and GST-PPBD$^{Mut}$ Protein Purification*

*E.coli* BL21(DE3) competent cells transformed with pGEX-6P2-GST, pGEX-6P2-GST-PPBD, and pGEX-6P2-GST-PPBD$^{Mut}$ respectively were incubated as the primary culture in Luria Bertani (LB) broth containing 100 ug/ml ampicillin overnight at 37℃. The culture was diluted into 500 mL LB broth and incubated at 37℃ till $OD_{600}$ reached 0.8. Protein expression was induced with 1 mM IPTG for 4 hours at 37℃ and shaking at 180 rpm. The cells were pelleted by centrifugation at 3000g for five minutes at 4℃. The pellet was resuspended in 50 ml buffer A (20 mM HEPES-KOH pH 6.8, 100 mM NaCl, 2 mM EDTA, 5 mM DTT (freshly added)) and lysed by sonication (15 minutes, output voltage 20, output control 30, duty cycle 30/40) using Branson Sonifier. The lysate was centrifuged at 14000g for 10 minutes at 4℃, and the supernatant was mixed with TritonX-100 to a final concentration of 0.1%. The lysate was incubated with 250 µl of a 1:1 slurry of glutathione beads (pre-equilibrated with buffer B (buffer A plus 1% Triton X-100)) at 4℃ for 2.5 hours, and beads were pelleted at 400g for three minutes at 4℃. The beads were washed thrice with buffer C (20 mM HEPES-KOH pH 6.8, 500 mM NaCl, 2 mM EDTA, 1% Triton X-100) and again thrice with buffer B followed by one wash with PBS, with centrifugation at 400g for three minutes at 4°C at each wash step. The protein was eluted from the beads with 200µl elution buffer (50 mM Tris-Cl pH 8.0, 50 mM reduced glutathione, 12 µl of 1M NaOH) and incubated overnight on a rotor at 4℃. Eluted protein in the supernatant was recovered by centrifugation at 400g at 4℃ and stored in aliquots at -80°C.

**Microscale Thermophoresis:**

2.5nM polyP$_{100}$-2X FITC was titrated against sixteen serially diluted concentrations of each of the proteins GST, GST-PPBD and GST-PPBD$^{Mut}$, starting from an initial protein concentration of 5μM. Microscale Thermophoresis was performed in Instrument Nano Temper, and the normalised fluorescence units ($F_{norm}$) and the fraction of polyP$_{100}$-2X FITC bound were plotted. From the normalised fluorescence units, the binding affinity of GST, GST-PPBD and GST-PPBD$^{Mut}$ against polyP were determined.

**Embryo collection:**

Flies acclimatised on the grape juice plates for 24 hours were used for egg laying for two hours on a fresh grape juice plate. Every two hours after egg-laying, we collected the embryos for polyP extraction. The eggs were incubated at 25℃ for 18 hours. We used 150 embryos for each set in our experiment.

**RNA extraction and qRT-PCR**

RNA isolation using TRIzol (Ambion life tech—15596018) method. cDNA conversion for 1μg of RNA was carried out using a cDNA conversion kit (Thermo Fisher—4368814). qPCR was carried out in 96 well plates in three technical replicates for each of the three biological replicate sets. Each biological replicate set had five time-matched non-feeding wandering third instar larvae.

**Fly lines:**

| **Name** | **Flybase ID** | **References** |
|---|---|---|
| *Canton-S* | FBsn0000274 | |
| *Tub-GAL4* | FBtp0002651 | |
| *Cg-GAL4* | FBtp0012452 | (3) |
| *Hml-GAL4 > UAS-GFP* | FBst0030140 | (4) |
| *Lz-GAL4 > UAS-mCD8-GFP* | FBst0006314 | (5) |
| *NosGAL4 VP16* | FBti0131634 | (6) |
| *y ; AttP40* | FBti0114379 | (7) |
| Cyto-FLYX: *w[*] ; p(pUAST-HA-ScPpx1)AttP40 / Cyo* | This work | |
| Nuc-FLYX: *w[*] ; p(pUAST-3X_Nuc-HA-ScPpx1)AttP40 / CyO* | This work | |
| ER-FLYX: *w[*] ;; p(pUAST-ER-HA-ScPpx1)AttP2 / TM3,Sb* | This work | |
| Mito-FLYX: *w[*] ;; p(pUAST-Mito-HA-ScPpx1)AttP2 / TM3,Sb* | This work | |
| pCyto-FLYX: *w[*] ; p(pUASp-HA-ScPpx1) / CyO* | This work | |

**Chemicals:**

| # | **Name** | **Company** | **Catalogue ID** |
|---|---|---|---|
| 1 | 100% molecular grade ethanol (diluent for DNA extraction) | Himedia | MB228 |
| 2 | 4% paraformaldehyde | Himedia | TCL119 |
| 3 | Acid phenol | Himedia | MB081 |
| 4 | Ammonium Molybdate | Himedia | GRM307 |
| 5 | BL21 DE3 competent cells | Himedia | MBT143 |
| 6 | Boric acid | Himedia | MB007 |

| | | | |
|---|---|---|---|
| 7 | Bovine Serum Albumin | Himedia | MB083 |
| 8 | Cell Mask Deep Red | Invitrogen | C10046 |
| 9 | Chloroform | Himedia | MB019 |
| 10 | DAPI hydrochloride | Himedia | MB097 |
| 11 | DL-DTT | Himedia | MB070 |
| 12 | EDTA | Himedia | GRM1370 |
| 13 | Goat anti-mouse-alexa555 | Invitrogen | A32727 |
| 14 | Goat anti-rabbit-alexa-488 | Invitrogen | A11034 |
| 15 | HEPES | Himedia | MB016 |
| 16 | HPLC grade water | Himedia | AS077 |
| 17 | Imidazole | Himedia | MB019 |
| 18 | L glutathione reduced | Sigma | G4251 |
| 19 | Lauryl sulphate (SDS) | Himedia | MB010 |
| 20 | Lithium chloride | Himedia | MB038 |
| 21 | Luria Broth | Himedia | M575 |
| 22 | Luria Broth Agar | Himedia | M1151 |
| 23 | Malachite Green Carbinol Hydrochloride | Glentham Life Sciences | GD8088 |
| 24 | Mouse anti-HA | Cell Signaling Technology | 2367 |
| 25 | Mouse anti-GST | Cell Signaling Technology | 2624 |
| 26 | Mouse anti-KDEL | Abcam | ab12223 |
| 27 | Mouse anti-ATP5A | Abcam | ab14748 |
| 28 | Ni-NTA agarose | Qiagen | 30210 |
| 29 | $PolyP_{14}$, $PolyP_{65}$, $PolyP_{130}$ (Ex-polyP set) | Regenetiss | Gifted by T.Shiba |

| | | | |
|---|---|---|---|
| 30 | Pierce glutathione agarose | Thermo scientific | 16100 |
| 31 | Potassium phosphate dibasic anhydrous | Himedia | MB044 |
| 32 | Rabbit Anti-Fibrillarin | CST | 2639 |
| 33 | Rabbit Anti-HA | CST | 3724S |
| 34 | RNase A | Himedia | DS0003 |
| 35 | Sodium chloride | Himedia | MB023 |
| 36 | Sodium hydroxide | Himedia | MB095 |
| 37 | Tris free base | Himedia | MB029 |
| 38 | Triton X100 | Himedia | MB031 |

| **Serial number** | **Plasmid name** | **Source** |
|---|---|---|
| 1 | pTrc-HisB-*ScPpx1* | Gifted by Adolfo Saiardi (*ScPpx1* gene ID-SGD:S000001244) |
| 2 | pGEX-6P2-GST-PPBD | This work |
| 3 | pGEX-6P2-GST-PPBD 8Mut | This work |
| 4 | pGEX-6P2-GST | This work |

**qRT-PCR primers**

| **Gene** | **Forward Primer (5’-3’)** | **Reverse Primer (5’-3’)** | **References** |
|---|---|---|---|
| *ecd* | CTGGCGGAGTTCTTAGATCG | GCATGGAGGGATTCTTCTTG | (8) |
| *ImPL2* | GCCGATACCTTCGTGTATCC | TTTCCGTCGTCAATCCAATAG | (9) |
| *Eip74EF* | TCCACAATCTGCTTAGCGGC | GACTGGGCGGAAATGAACCT | (10) |
| *inR* | AACAGTGGCGGATTCGGTT | TACTCGGAGCATTGGAGGCAT | (11) |
| *chico* | GGCATACGGGCAGCTAGAC | TTCTTGAGGTAGCCACTCAGC | (11) |

| | | | |
|---|---|---|---|
| *dfoxo* | TCGAGTGCAATGTCGAGGAG | AGCGGTATATTGATGTCCAGCAG | (11) |
| *dilp2* | ATGGTGTGCGAGGAGTATAATCC | TCGGCACCGGGCATG | (12) |
| *s6k* | TGACCTAGAACCGGAATTGTG | TCCTCGCAGAGCTGTATGG | (11) |
| *4ebp* | CACTCCTGGAGGCACCA | GAGTTCCCCTCAGCAAGCAA | (12) |
| *tor* | GCTCAGAGGCGAGAGACAAG | CCAGCTCACGGAGGATAAAG | (11) |
| *rp49* | TCCTACCAGCTTCAAGATGAC | CACGTTGTGCACCAGGAACT | This work |

**Sequence of codon optimised *HA-ScPpx1* for creation of Cyto-FLYX system**

ATGTATCCGTATGATGTTCCGGATTATGCATCGCCTTTGAGAAAGACGGTTCCTGAAT

TTTTGGCACACTTAAAATCACTGCCAATTTCAAAAATTGCAAGCAACGATGTGTTAA

CAATATGTGTTGGTAACGAGTCAGCAGATATGGACTCAATTGCTAGTGCAATCACTTA

TTCGTACTGCCAATACATATATAATGAAGGTACTTACTCGGAGGAGAAAAAGAAAGG

AAGCTTTATTGTCCCAATTATCGACATTCCTAGAGAAGACCTCAGTTTAAGAAGAGA

CGTAATGTATGTCCTAGAAAAACTGAAAATTAAGGAGGAGGAATTATTCTTCATTGAA

GATTTAAAGAGCTTGAAGCAAAATGTCTCCCAGGGTACTGAATTAAACTCTTACTTG

GTAGATAATAACGATACACCAAAGAATTTGAAAAATTATATAGATAACGTCGTTGGCA

TTATAGACCACCATTTTGACTTGCAAAAACATTTGGATGCTGAACCTCGGATTGTAAA

AGTGTCCGGCAGTTGCTCATCGCTGGTTTTTAACTACTGGTATGAAAAATTGCAAGG

TGACCGTGAAGTGGTGATGAACATTGCACCACTTTTGATGGGGGCCATCTTAATAGA

CACTTCAAATATGAGGCGCAAAGTCGAGGAAAGTGATAAATTAGCTATCGAGAGATG

CCAAGCTGTTCTTAGTGGTGCGGTTAATGAAGTGTCTGCGCAAGGTTTAGAGGACAG

CAGTGAGTTTTATAAAGAGATAAAATCAAGAAAGAACGATATTAAAGGATTTTCGGT

AAGCGATATTCTAAAGAAGGACTACAAACAATTCAATTTCCAAGGAAAGGGACACA

AAGGGTTAGAGATTGGTCTTTCATCAATAGTAAAAAGAATGTCTTGGCTATTCAATGA

ACACGGTGGTGAAGCAGATTTCGTCAACCAATGCAGAAGATTTCAGGCGGAGAGGG

GGCTCGATGTATTGGTTCTGTTGACTTCATGGAGGAAAGCTGGTGATTCACACAGAG

AATTGGTCATATTGGGAGACTCTAACGTGGTACGTGAACTCATTGAAAGGGTTAGCG

ACAAGCTCCAACTTCAATTATTTGGGGGCAATCTTGATGGAGGTGTGGCGATGTTTA

AGCAACTGAACGTCGAGGCCACCAGAAAGCAAGTCGTCCCCTATTTAGAGGAAGCG

TACTCAAACCTGGAAGAGTGA

**Sequence of codon optimised 3X-NLS-*HA-ScPpx1* for creation of Nuc-FLYX system**

ATGCCCAAGAAGAAGAGGAAGGTTCCCAAGAAGAAGAGGAAGGTTCCCAAGAAGA

AGAGGAAGGTTGGCGGCCGCTATCCGTATGATGTTCCGGATTATGCATCGCCTTTGAG

AAAGACGGTTCCTGAATTTTTGGCACACTTAAAATCACTGCCAATTTCAAAAATTGC

AAGCAACGATGTGTTAACAATATGTGTTGGTAACGAGTCAGCAGATATGGACTCAAT

TGCTAGTGCAATCACTTATTCGTACTGCCAATACATATATAATGAAGGTACTTACTCGG

AGGAGAAAAAGAAAGGAAGCTTTATTGTCCCAATTATCGACATTCCTAGAGAAGACC

TCAGTTTAAGAAGAGACGTAATGTATGTCCTAGAAAAACTGAAAATTAAGGAGGAG

GAATTATTCTTCATTGAAGATTTAAAGAGCTTGAAGCAAAATGTCTCCCAGGGTACTG

AATTAAACTCTTACTTGGTAGATAATAACGATACACCAAAGAATTTGAAAAATTATATA

GATAACGTCGTTGGCATTATAGACCACCATTTTGACTTGCAAAAACATTTGGATGCTG

AACCTCGGATTGTAAAAGTGTCCGGCAGTTGCTCATCGCTGGTTTTTAACTACTGGTA

TGAAAAATTGCAAGGTGACCGTGAAGTGGTGATGAACATTGCACCACTTTTGATGGG

GGCCATCTTAATAGACACTTCAAATATGAGGCGCAAAGTCGAGGAAAGTGATAAATT

AGCTATCGAGAGATGCCAAGCTGTTCTTAGTGGTGCGGTTAATGAAGTGTCTGCGCA

AGGTTTAGAGGACAGCAGTGAGTTTTATAAAGAGATAAAATCAAGAAAGAACGATAT

TAAAGGATTTTCGGTAAGCGATATTCTAAAGAAGGACTACAAACAATTCAATTTCCA

AGGAAAGGGACACAAAGGGTTAGAGATTGGTCTTTCATCAATAGTAAAAAGAATGT

CTTGGCTATTCAATGAACACGGTGGTGAAGCAGATTTCGTCAACCAATGCAGAAGAT

TTCAGGCGGAGAGGGGGCTCGATGTATTGGTTCTGTTGACTTCATGGAGGAAAGCTG

GTGATTCACACAGAGAATTGGTCATATTGGGAGACTCTAACGTGGTACGTGAACTCA

TTGAAAGGGTTAGCGACAAGCTCCAACTTCAATTATTTGGGGGCAATCTTGATGGAG

GTGTGGCGATGTTTAAGCAACTGAACGTCGAGGCCACCAGAAAGCAAGTCGTCCCC

TATTTAGAGGAAGCGTACTCAAACCTGGAAGAGTGA

**Sequence of codon optimised Mito-*HA-ScPpx1* for creation of Mito-FLYX system**

ATGTCCGTCCTGACGCCGCTGCTGCTGCGGGGCTTGACAGGCTCGGCCCGGCGGCT
CCCAGTGCCGCGCGCCAAGATCCATTCGTTTGCCGCCGGCGGCCGCTATCCGTATGAT
GTTCCGGATTATGCATCGCCTTTGAGAAAGACGGTTCCTGAATTTTTGGCACACTTAA
AATCACTGCCAATTTCAAAAATTGCAAGCAACGATGTGTTAACAATATGTGTTGGTAA
CGAGTCAGCAGATATGGACTCAATTGCTAGTGCAATCACTTATTCGTACTGCCAATAC
ATATATAATGAAGGTACTTACTCGGAGGAGAAAAAGAAAGGAAGCTTTATTGTCCCA
ATTATCGACATTCCTAGAGAAGACCTCAGTTTAAGAAGAGACGTAATGTATGTCCTAG
AAAAACTGAAAATTAAGGAGGAGGAATTATTCTTCATTGAAGATTTAAAGAGCTTGA
AGCAAAATGTCTCCCAGGGTACTGAATTAAACTCTTACTTGGTAGATAATAACGATAC
ACCAAAGAATTTGAAAAATTATATAGATAACGTCGTTGGCATTATAGACCACCATTTT
GACTTGCAAAAACATTTGGATGCTGAACCTCGGATTGTAAAAGTGTCCGGCAGTTGC
TCATCGCTGGTTTTTAACTACTGGTATGAAAAATTGCAAGGTGACCGTGAAGTGGTG
ATGAACATTGCACCACTTTTGATGGGGGCCATCTTAATAGACACTTCAAATATGAGGC
GCAAAGTCGAGGAAAGTGATAAATTAGCTATCGAGAGATGCCAAGCTGTTCTTAGTG
GTGCGGTTAATGAAGTGTCTGCGCAAGGTTTAGAGGACAGCAGTGAGTTTTATAAAG
AGATAAAATCAAGAAAGAACGATATTAAAGGATTTTCGGTAAGCGATATTCTAAAGA
AGGACTACAAACAATTCAATTTCCAAGGAAAGGGACACAAAGGGTTAGAGATTGGT
CTTTCATCAATAGTAAAAAGAATGTCTTGGCTATTCAATGAACACGGTGGTGAAGCA
GATTTCGTCAACCAATGCAGAAGATTTCAGGCGGAGAGGGGGCTCGATGTATTGGTT
CTGTTGACTTCATGGAGGAAAGCTGGTGATTCACACAGAGAATTGGTCATATTGGGA

GACTCTAACGTGGTACGTGAACTCATTGAAAGGGTTAGCGACAAGCTCCAACTTCA
ATTATTTGGGGGCAATCTTGATGGAGGTGTGGCGATGTTTAAGCAACTGAACGTCGA
GGCCACCAGAAAGCAAGTCGTCCCCTATTTAGAGGAAGCGTACTCAAACCTGGAAG
AGTGA

**Sequence of codon optimised ER-*HA-ScPpx1* for creation of ER-FLYX system**

ATGAGCTTTGTGAGCCTGCTGCTGGTGGGCATTCTGTTTTGGGCGACCGAAGCGGAA
CAGCTGACCAAATGCGAAGTGTTTTGCGGCCGCTATCCGTATGATGTTCCGGATTATG
CATCGCCTTTGAGAAAGACGGTTCCTGAATTTTTGGCACACTTAAAATCACTGCCAA
TTTCAAAAATTGCAAGCAACGATGTGTTAACAATATGTGTTGGTAACGAGTCAGCAG
ATATGGACTCAATTGCTAGTGCAATCACTTATTCGTACTGCCAATACATATATAATGAA
GGTACTTACTCGGAGGAGAAAAAGAAAGGAAGCTTTATTGTCCCAATTATCGACATT
CCTAGAGAAGACCTCAGTTTAAGAAGAGACGTAATGTATGTCCTAGAAAAACTGAA
AATTAAGGAGGAGGAATTATTCTTCATTGAAGATTTAAAGAGCTTGAAGCAAAATGT
CTCCCAGGGTACTGAATTAAACTCTTACTTGGTAGATAATAACGATACACCAAAGAAT
TTGAAAAATTATATAGATAACGTCGTTGGCATTATAGACCACCATTTTGACTTGCAAA
AACATTTGGATGCTGAACCTCGGATTGTAAAAGTGTCCGGCAGTTGCTCATCGCTGG
TTTTTAACTACTGGTATGAAAAATTGCAAGGTGACCGTGAAGTGGTGATGAACATTG
CACCACTTTTGATGGGGGCCATCTTAATAGACACTTCAAATATGAGGCGCAAAGTCG
AGGAAAGTGATAAATTAGCTATCGAGAGATGCCAAGCTGTTCTTAGTGGTGCGGTTA
ATGAAGTGTCTGCGCAAGGTTTAGAGGACAGCAGTGAGTTTTATAAAGAGATAAAAT
CAAGAAAGAACGATATTAAAGGATTTTCGGTAAGCGATATTCTAAAGAAGGACTACA
AACAATTCAATTTCCAAGGAAAGGGACACAAAGGGTTAGAGATTGGTCTTTCATCAA

TAGTAAAAAGAATGTCTTGGCTATTCAATGAACACGGTGGTGAAGCAGATTTCGTCA

ACCAATGCAGAAGATTTCAGGCGGAGAGGGGGCTCGATGTATTGGTTCTGTTGACTT

CATGGAGGAAAGCTGGTGATTCACACAGAGAATTGGTCATATTGGGAGACTCTAACG

TGGTACGTGAACTCATTGAAAGGGTTAGCGACAAGCTCCAACTTCAATTATTTGGGG

GCAATCTTGATGGAGGTGTGGCGATGTTTAAGCAACTGAACGTCGAGGCCACCAGA

AAGCAAGTCGTCCCCTATTTAGAGGAAGCGTACTCAAACCTGGAAGAGTGA

**Sequence of GST-PPBD**

ATGTCCCCTATACTAGGTTATTGGAAAATTAAGGGCCTTGTGCAACCCACTCGACTTC
TTTTGGAATATCTTGAAGAAAAATATGAAGAGCATTTGTATGAGCGCGATGAAGGTG
ATAAATGGCGAAACAAAAAGTTTGAATTGGGTTTGGAGTTTCCCAATCTTCCTTATTA
TATTGATGGTGATGTTAAATTAACACAGTCTATGGCCATCATACGTTATATAGCTGACA
AGCACAACATGTTGGGTGGTTGTCCAAAAGAGCGTGCAGAGATTTCAATGCTTGAA
GGAGCGGTTTTGGATATTAGATACGGTGTTTCGAGAATTGCATATAGTAAAGACTTTG
AAACTCTCAAAGTTGATTTTCTTAGCAAGCTACCTGAAATGCTGAAAATGTTCGAAG
ATCGTTTATGTCATAAAACATATTTAAATGGTGATCATGTAACCCATCCTGACTTCATG
TTGTATGACGCTCTTGATGTTGTTTTATACATGGACCCAATGTGCCTGGATGCGTTCCC
AAAATTAGTTTGTTTTAAAAAACGTATTGAAGCTATCCCACAAATTGATAAGTACTTG
AAATCCAGCAAGTATATAGCATGGCCTTTGCAGGGCTGGCAAGCCACGTTTGGTGGT
GGCGACCATCCTCCAAAATCGGATCTGGAAGTTCTGTTCCAGGGGCCCCTGGGATCC
ATGGATCTGTATGATGATGATGATAAAGAATTCATGGAAGGACGTTTCCGTCATCAGG
ATGTGCGTAGTCGCACCGCCAGCAGCCTCGCCAACCAGTATCACATCGACAGCGAAC
AGGCCCGACGAGTGCTGGATACCACTATGCAAATGTACGAACAGTGGCGGGAACAG
CAACCGAAGCTGGCGCATCCGCAACTGGAGGCGCTACTGCGATGGGCCGCCATGCT
GCATGAGGTCGGGTTGAATATCAACCACAGCGGTTTGCATCGCCACTCCGCTTATATT
CTGCAAAACAGTGACTTGCCGGGTTTTAATCAGGAACAGCAGCTGATGATGGCGAC
ACTGGTGCGCTATCACCGTAAAGCGATTAAGCTCGACGATCTGCCGCGCTTTACCTTG
TTTAAGAAGAAACAGTTCCTGCCACTGATACAGCTATTGCGCCTTGGCGTATTACTAA
ACAATCAACGTCAGGCAACCACCACACCGCCAACATTGACACTGATTACTGATGACA
GTCACTGGACACTGCGTTTCCCGCATGACTGGTTTAGTCAGAATGCGCTGGTACTGC

TTGATCTGGAAAAGGAGCAAGAATACTGGGAAGGCGTGGCTGGCTGGCGGTTGAAA

ATTGAAGAAGAAAGTACACCTGAAATCGCCGCTTAA

**Protein sequence with Histidines, Arginines, and Lysines that are predicted for polyP binding are underlined**

MSPILGYWKIKGLVQPTRLLLEYLEEKYEEHLYERDEGDKWRNKKFELGLEFPNLPYYI

DGDVKLTQSMAIIRYIADKHNMLGGCPKERAEISMLEGAVLDIRYGVSRIAYSKDFETLK

VDFLSKLPEMLKMFEDRLCHKTYLNGDHVTHPDFMLYDALDVVLYMDPMCLDAFPKL

VCFKKRIEAIPQIDKYLKSSKYIAWPLQGWQATFGGGDHPPKSDLEVLFQGPLGSMDLY

DDDDKEFMEGRFRHQDVRSRTASSLANQYHIDSEQARRVLDTTMQMYEQWREQQPKL

AHPQLEALLRWAAMLHEVGLNINHSGLHRHSAYILQNSDLPGFNQEQQLMMATLVRYH

RKAIKLDDLPRFTLFKKKQFLPLIQLLRLGVLLNNQRQATTTPPTLTLITDDSHWTLRFP

HDWFSQNALVLLDLEKEQEYWEGVAGWRLKIEEESTPEIAA*

**Sequence of GST-PPBD$^{Mut}$**

ATGTCCCCTATACTAGGTTATTGGAAAATTAAGGGCCTTGTGCAACCCACTCGACTTC
TTTTGGAATATCTTGAAGAAAAATATGAAGAGCATTTGTATGAGCGCGATGAAGGTG
ATAAATGGCGAAACAAAAAGTTTGAATTGGGTTTGGAGTTTCCCAATCTTCCTTATTA
TATTGATGGTGATGTTAAATTAACACAGTCTATGGCCATCATACGTTATATAGCTGACA
AGCACAACATGTTGGGTGGTTGTCCAAAAGAGCGTGCAGAGATTTCAATGCTTGAA
GGAGCGGTTTTGGATATTAGATACGGTGTTTCGAGAATTGCATATAGTAAAGACTTTG
AAACTCTCAAAGTTGATTTTCTTAGCAAGCTACCTGAAATGCTGAAAATGTTCGAAG
ATCGTTTATGTCATAAAACATATTTAAATGGTGATCATGTAACCCATCCTGACTTCATG
TTGTATGACGCTCTTGATGTTGTTTTATACATGGACCCAATGTGCCTGGATGCGTTCCC
AAAATTAGTTTGTTTTAAAAAACGTATTGAAGCTATCCCACAAATTGATAAGTACTTG
AAATCCAGCAAGTATATAGCATGGCCTTTGCAGGGCTGGCAAGCCACGTTTGGTGGT
GGCGACCATCCTCCAAAATCGGATCTGGAAGTTCTGTTCCAGGGGCCCCTGGGATCC
ATGGATCTGTATGATGATGATGATAAAGAATTCATGGAAGGACGGTTCCGCCATCAGG
ATGTGCGATCACGCACAGCGTCTTCGCTCGCCAACCAGTACCACATTGATTCCGAAC
AAGCCCGCCGCGTTTTGGACACGACTATGCAAATGTACGAACAGTGGCGGGAGCAG
CAGCCCAAGTTGGCTCACCCTCAGCTAGAAGCACTTCTAAGGTGGGCGGCCATGTTG
CACGAGGTAGGCCTCAATATTAATGCAAGTGGTCTGGCTGCAGCCTCGGCATATATCC
TTCAAAACAGCGACCTGCCGGGATTTAATCAGGAGCAACAGCTCATGATGGCTACGC
TAGTGCGCTACGCCGCCGCCGCCATTGCGCTGGACGACCTCCCACGTTTCACCTTGT
TCAAGAAAAAGCAGTTTCTGCCACTGATACAATTACTGAGACTGGGGGTGCTCTTGA
ATAACCAGCGTCAGGCCACTACCACGCCGCCCACATTGACACTGATCACCGATGATT
CCCATTGGACCCTGCGATTTCCCCACGATTGGTTCAGCCAGAACGCCCTGGTCCTTC

TGGACCTGGAGAAGGAACAGGAGTATTGGGAGGGCGTCGCGGGCTGGAGGCTGAA

AATCGAGGAGGAGAGCACCCCGGAGATCGCGGCTTAA

**Protein sequence with Alanines mutation (underlined):**

MSPILGYWKIKGLVQPTRLLLEYLEEKYEEHLYERDEGDKWRNKKFELGLEFPNLPYYI

DGDVKLTQSMAIIRYIADKHNMLGGCPKERAEISMLEGAVLDIRYGVSRIAYSKDFETLK

VDFLSKLPEMLKMFEDRLCHKTYLNGDHVTHPDFMLYDALDVVLYMDPMCLDAFPKL

VCFKKRIEAIPQIDKYLKSSKYIAWPLQGWQATFGGGDHPPKSDLEVLFQGPLGSMDLY

DDDDKEFMEGRFRHQDVRSRTASSLANQYHIDSEQARRVLDTTMQMYEQWREQQPKL

AHPQLEALLRWAAMLHEVGLNINASGLAAASAYILQNSDLPGFNQEQQLMMATLVRY

AAAAIALDDLPRFTLFKKKQFLPLIQLLRLGVLLNNQRQATTTPPTLTLITDDSHWTLRF

PHDWFSQNALVLLDLEKEQEYWEGVAGWRLKIEEESTPEIAA*

**Figure legends:**

**Supplementary Fig 1: A)** Schematic of citrate-saturated phenol-chloroform-based polyphosphate extraction from *Drosophila* larvae. **B)** Schematic of polyphosphate quantification using Malachite Green.

**Supplementary Fig 2: GST::PPBD staining in fly tissues. A)** GST::PPBD staining using different fixatives. Salivary glands staining with GST-PPBD and GST control in Bouin's fixative. Salivary glands stained with GST-PPBD and GST control in methanol fixative. Staining of larval salivary glands with DAPI (nucleus; cyan) and anti-GST antibody (GST; orange hot). The greyscale panel represents the GST channel. Scale bar - 20μm. *Drosophila* strain used - *CantonS*.

**Supplementary Fig 3: Polyphosphate staining of several *Drosophila* tissues with PPBD. (A, C, E, G, I, K, M, O)** Samples were incubated with GST-PPBD and stained with DAPI (nucleus; cyan) and anti-GST antibody (orange hot or greyscale). **B, D, F, H, J, L**, **N, P)** Negative control: tissues were incubated with GST protein and stained with DAPI (nucleus; cyan) and anti-GST antibody (GST; orange hot or greyscale). Scale bar - 20μm. *Drosophila* strain used - *CantonS*.

**Supplementary Fig 4: Polyphosphate affects hemolymph clotting. A-E)** Depiction of hemolymph clotting revealed by fibre formation at the edge of the hemolymph drop. **B-C)** Control (*AttP40* strain) (B) showing enriched with clot fibres and *hml* mutant (C) with minor clot fibres at the edges of the drop. **D-E)** Quantification of hemolymph clot fibre number (D) and length (E) in Control and *hml* mutants. **F-V)** Hemolymph clotting analysis followed by *ex vivo* polyP treatment. The relative number of clot fibres formed at the edge of the hemolymph drop incubated with **F)** water, N=5, **G)** Pi, N=5, **H)** PolyP$_{14,}$ N=4, **I)** PolyP$_{65}$, N=5, **J)** with PolyP$_{130,}$ N=4. **K-R)** Quantifications of fibre numbers and length in F-J. Statistics: Student's *t*-test with $p > 0.001$, error bar - s.e.m. **S-V)** Hemolymph clot fibre branching analysis. Fold change of clot fibre branches in F-J. Statistics: Student's *t*-test with $p > 0.001$, error bar - s.e.m. Scale bar for (J-N)- 500 px, Image dimensions in pixel: 2688 x 2200.

**Supplementary Fig 5: (A-E)** Clot fibre formation and quantification of fibre phenotype upon 30 minutes incubation with water (A), Pi (B), P$_{14}$ C), P$_{65}$ (D), P$_{130}$ (E). Scale bar for (A-E)- 500 pixels. Image dimensions in pixels: 3513 x 2712. Statistics: Student's *t*-test with $p > 0.001$, error bar - s.e.m. **(F-I)** Clot fibre formation and quantification of fibre phenotype upon 30 minutes incubation with (F) water, (G) P$_{65}$ (250µM Pi terms), (H) P$_{65}$ (125µM Pi terms). Statistics: Student's t-test with p > 0.001, error bar - s.e.m.

**Supplementary Fig 6: Phenotypic analysis of Cyto-FLYX w.r.t control A)** Cyto-FLYX does not change larval weight, where sets of 10 wandering third instar larvae taken from bottle are ethanol and water washed and weighed in a microcentrifuge tube. m=10, N=5. **B)** Cyto-FLYX does not change larval length, where wandering third instar larvae taken from bottles are ethanol and water washed, kept on a slide and imaged, N=29. **C)** Cyto-FLYX does not change adult fly weight, where sets of 10 5-day old adult flies taken from bottles are weighed in a microcentrifuge tube. m=10, N=6. **D)** Cyto-FLYX does not change pupal size (length) , where >65 mid pupa (1day old pupa) taken from bottles are kept on a slide and imaged, N=30. **(E-F)** Cyto-FLYX does not change fly fecundity. **E)** Expression of pUASp-Cyto-FLYX driven by *NosGAL4* in adult ovaries (3 day old flies) checked by anti-HA staining in magenta showing expression in ovarian stages. **F)** Cyto-FLYX does not show any difference in the number of eggs laid per fly every 24 hours w.r.t control flies. 10 sets of five females mated with males for two days were scored for next 24 hours after a 12 hour of acclimatization. Student's *t*-test with $p >$ 0.001, error bar - s.e.m.

**Supplementary Fig 7: Transcriptomic analysis of *TubGAL4* driven Cyto-FLYX w.r.t. control A)** qRT-PCR of RNA extracted from third instar non-feeding wandering third instar larvae shows no change in TOR, InR, Ecd related major transcripts between Cyto-FLYX and control. **B)** Principal Component Analysis of two biological replicates of samples- Cyto-FLYX and control, both driven by *TubGAL4*.

**Supplementary Fig 8: Transcriptomic analysis of *TubGAL4* driven Cyto-FLYX w.r.t. Control**, **GSEA and heat map**

**(A-C)** GO analysis in Cyto-FLYX and control larvae showing Biological processes (A), Cellular components (B), and Molecular functions (C), **D)** Heatmap of three gene sets enriched in Cyto-FLYX versus control larval RNA samples from the GO analysis - showing genes from mitochondrial translation, cytoplasmic translation, and ribosome biogenesis processes.

**Tables:**

**Table 1: Absolute quantification of polyP levels**

| | **PolyP quantification** |
|---|---|
| Normalised to protein content | 46.80 ± 3.95 picomoles in Pi terms / mg protein |
| Normalised to number of larvae | 419.3 ± 36.83 picomoles in Pi terms / larva |

**Table 2: PolyP quantification from embryo (Fig 1C)**

| **Embryo stages (hours)** | **polyP (pmoles of Pi/embryo)** |
|---|---|
| 1-3 hrs | 7.961 ± 3.27 |
| 3-5 hrs | 8.825 ± 3.57 |
| 5-7 hrs | 7.135 ± 1.39 |
| 7-9 hrs | 6.930 ± 1.48 |
| 9-11 hrs | 22.47 ± 13.22 |
| 12-14 hrs | 14.57 ± 4.49 |
| 14-16 hrs | 10.26 ± 3.78 |
| 16-18 hrs | 13.67 ± 3.38 |

**Table 3: PolyP quantification across fly life cycle (Fig 1D)**

| **Developmental stages** | **polyP (pmoles of Pi/mg protein)** |
|---|---|
| Embryo | 2.808 ± 1.0 |
| First instar | 1.714 ± 0.94 |
| Second instar | 2.038 ± 1.0 |
| Feeding third instar | 1.002 ± 0.53 |

| | |
|---|---|
| Non-feeding third instar (wandering) | 44.06 ± 7.04 |
| Prepupa | 15.63 ± 1.34 |
| One day post-pupariation | 16.88 ± 3.93 |
| Pharate | 1.398 ± 0.24 |
| Three days post-eclosion | 2.445 ± 0.78 |
