## Supplementary Figures for "Probing metazoan polyphosphate biology using *Drosophila* reveals novel and conserved polyP functions"

**Supplementary 1**

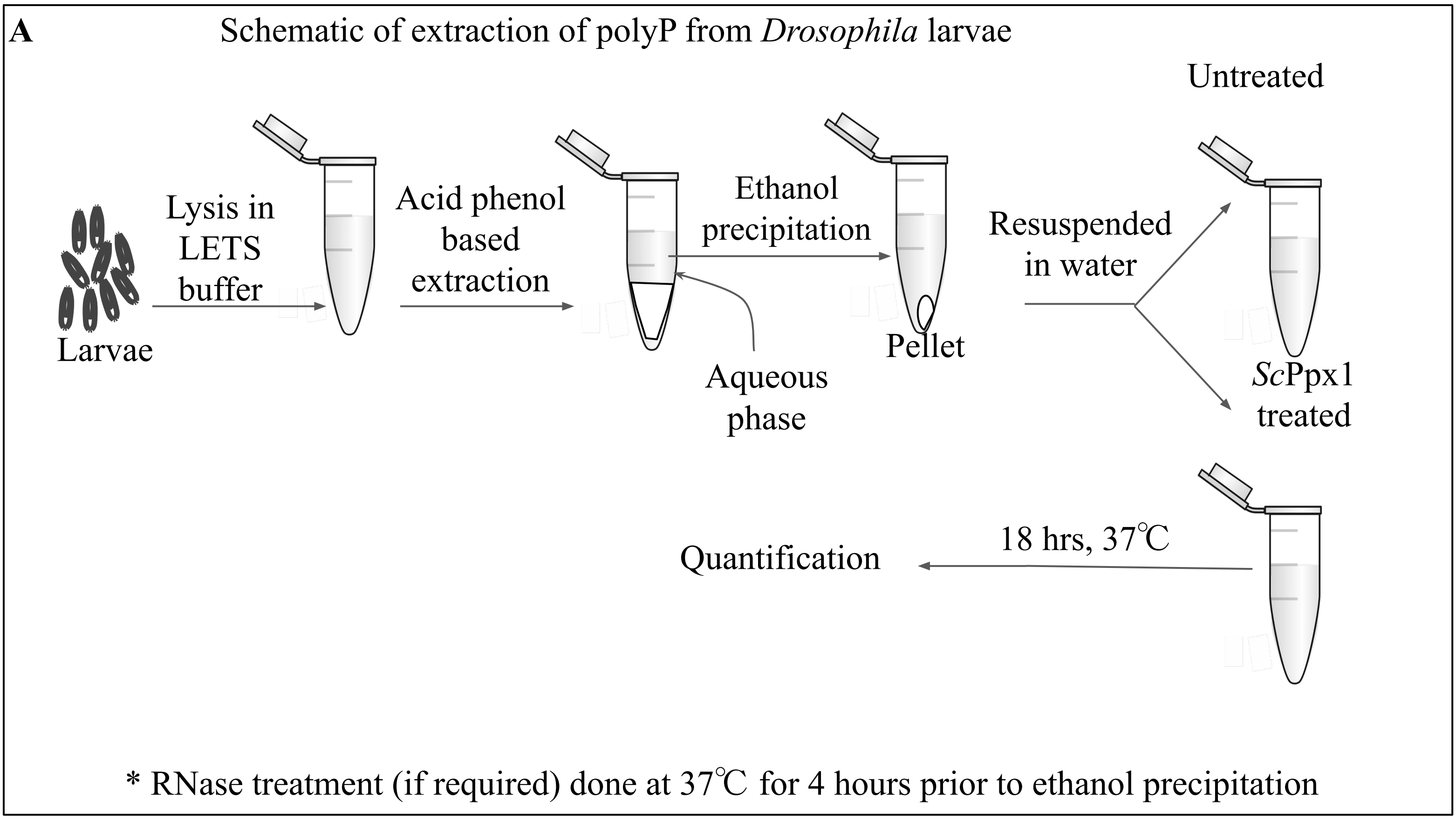


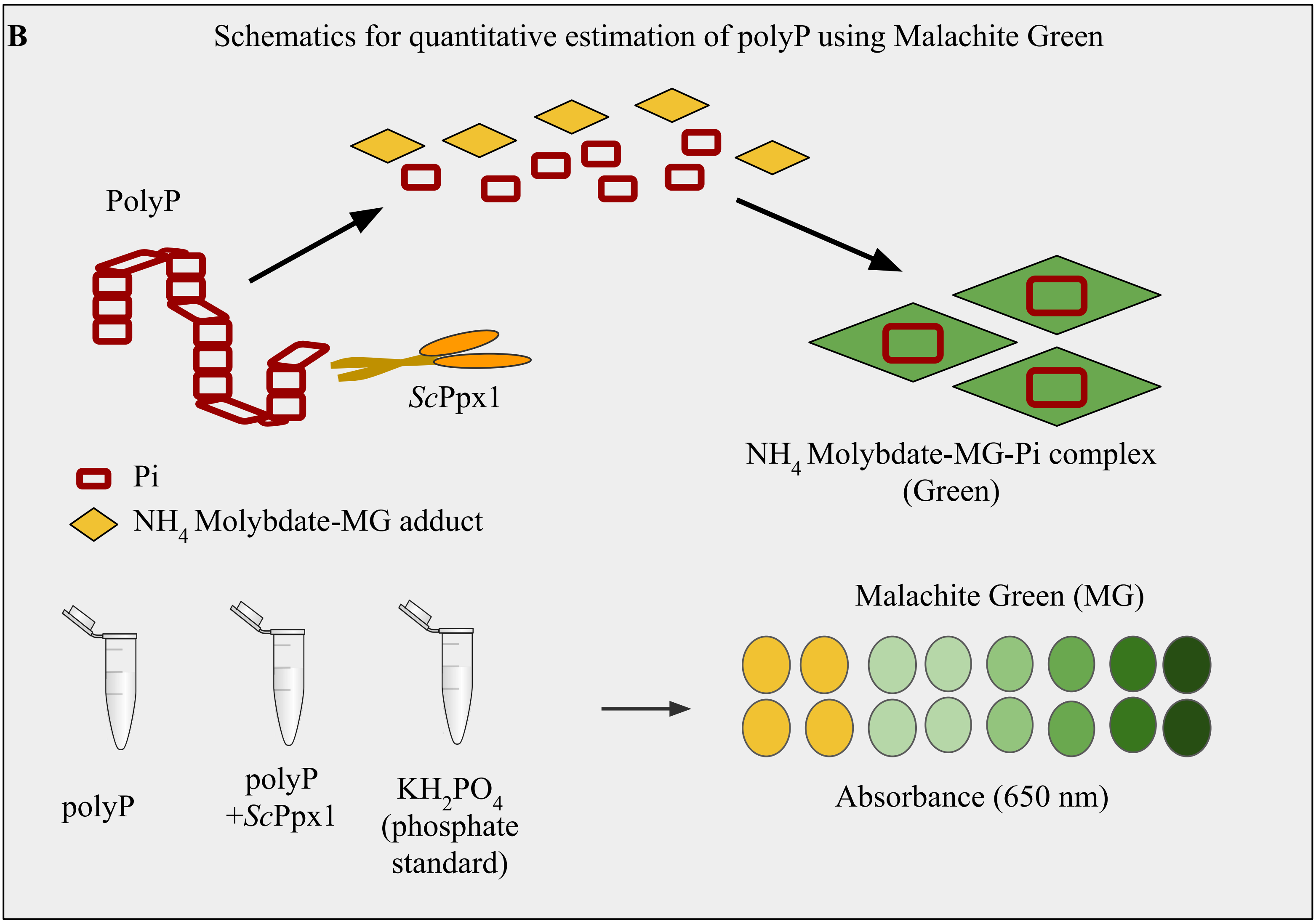

### Supplementary 2

**A** GST::PPBD staining in salivary glands with different fixatives

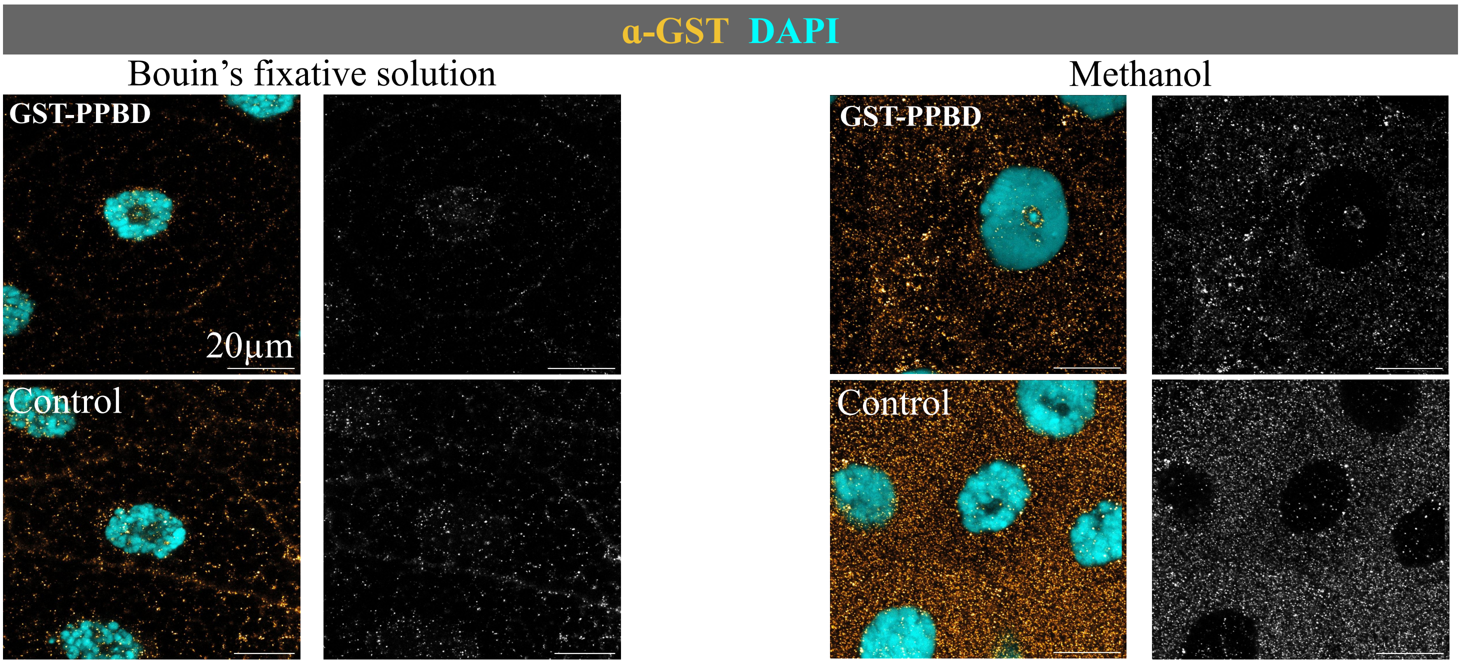

#### Supplementary 3

GST-PPBD staining in different larval tissues

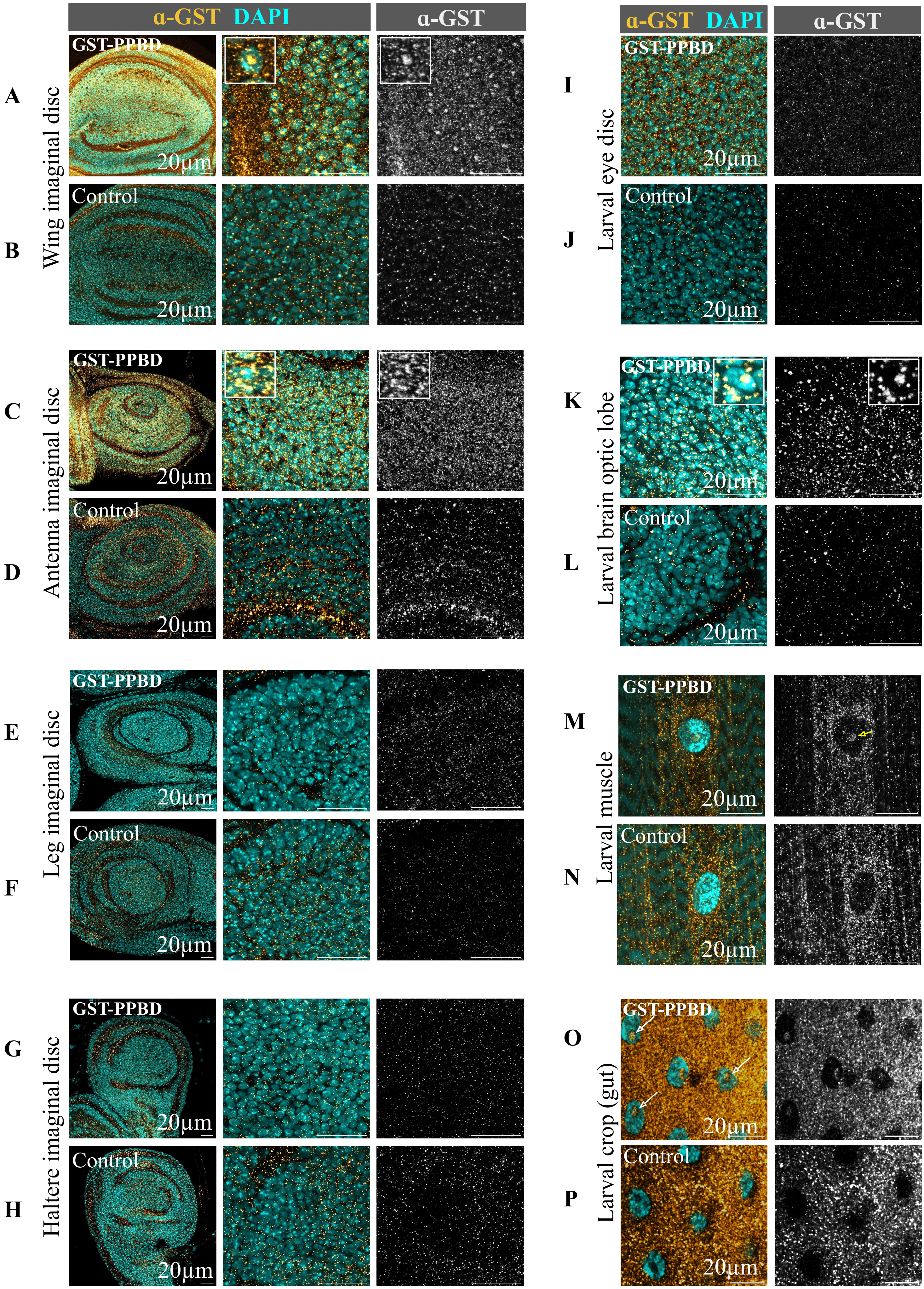

**Supplementary 4**

PolyP increases hemolymph clot fibre number density

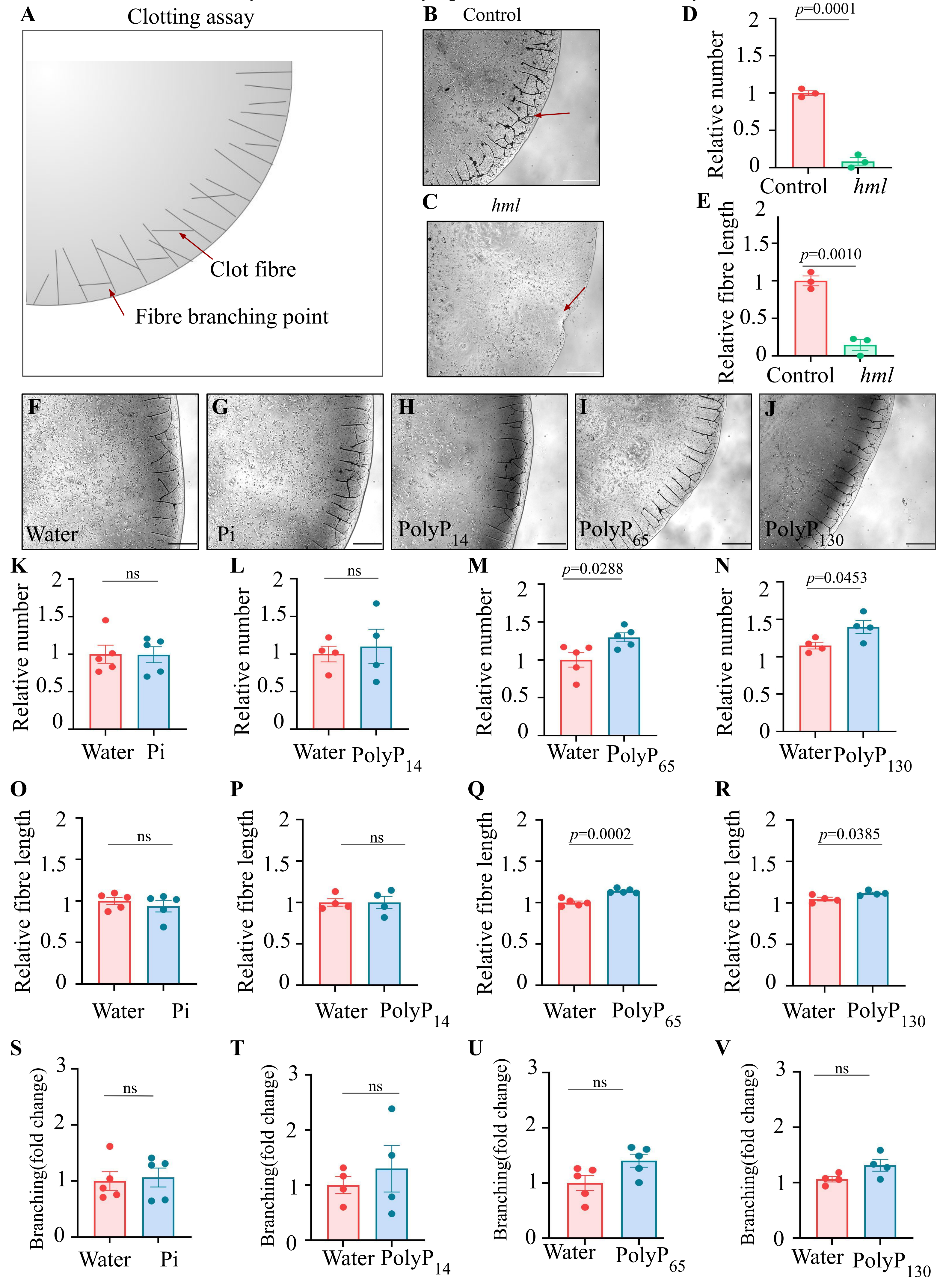

**Supplementary 5**

Hemolymph clot fibre formation across edge of drop after 30 minutes of incubation

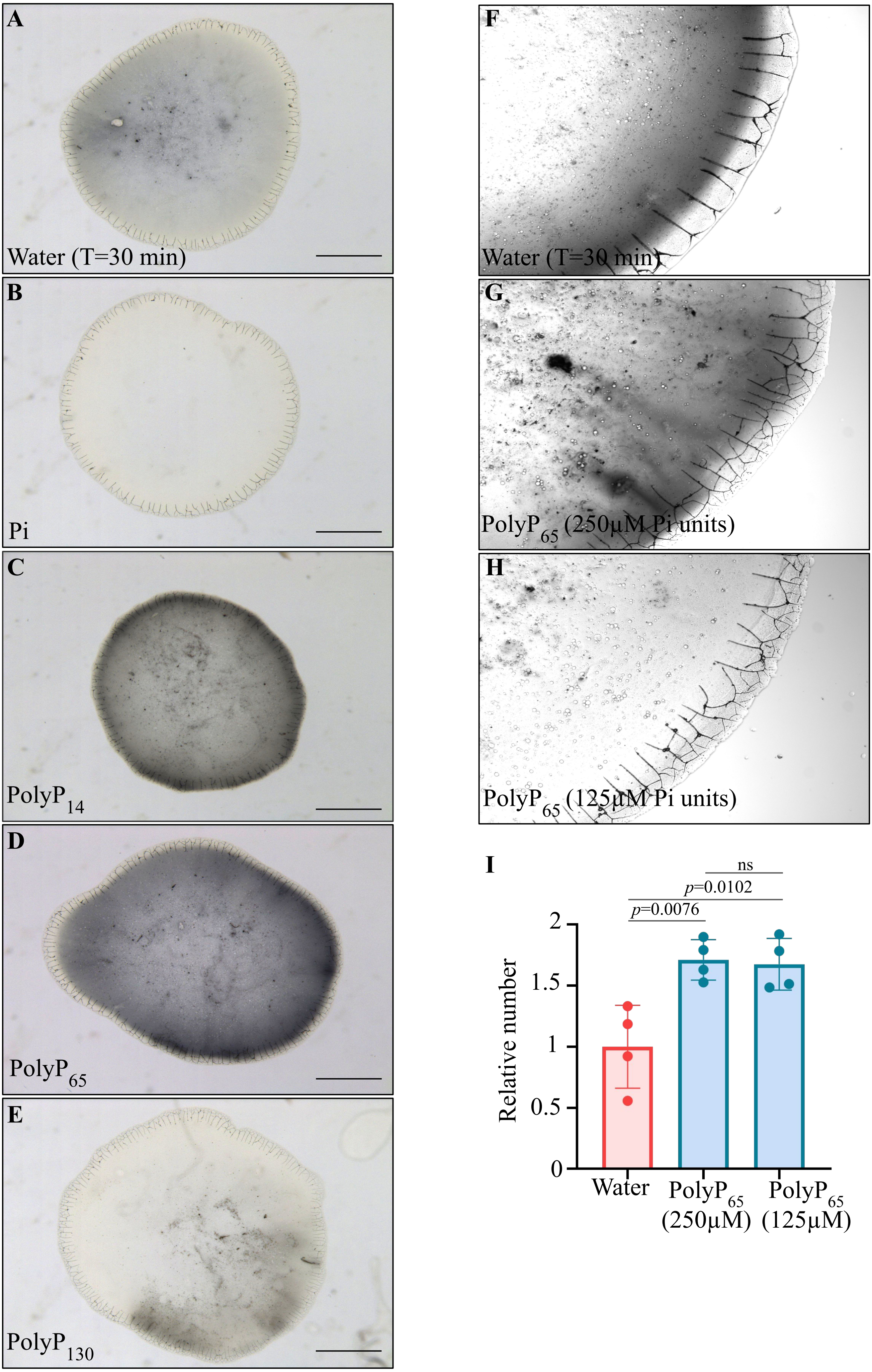

**Supplementary 6**

#### Cyto-FLYX does not change larval weight and length

A

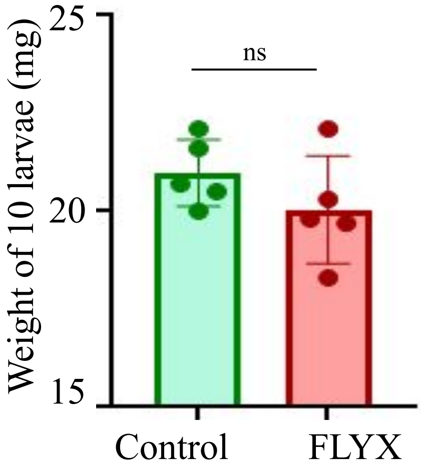


B

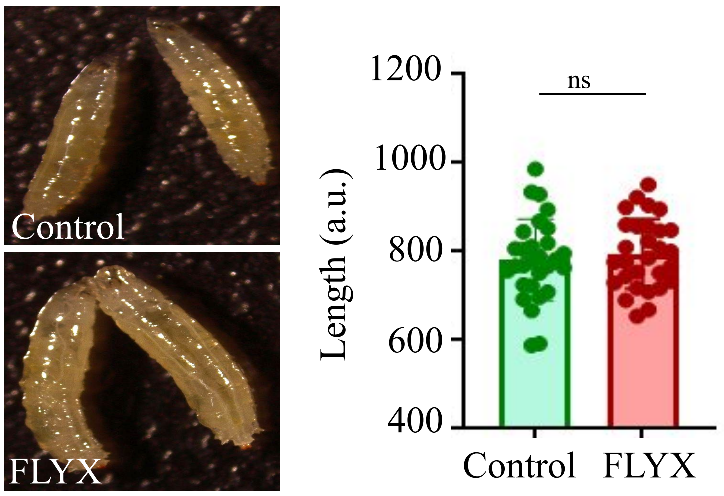


#### Cyto-FLYX does not change adult fly weight and length

C

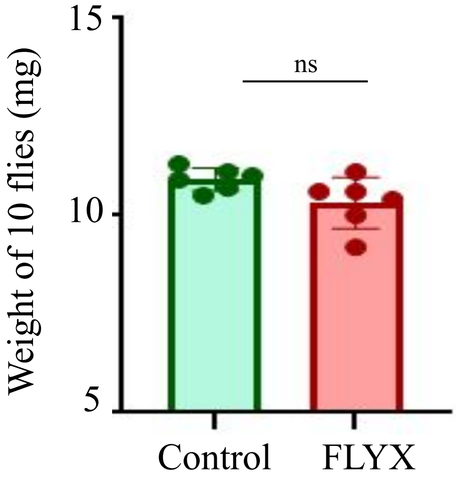


D

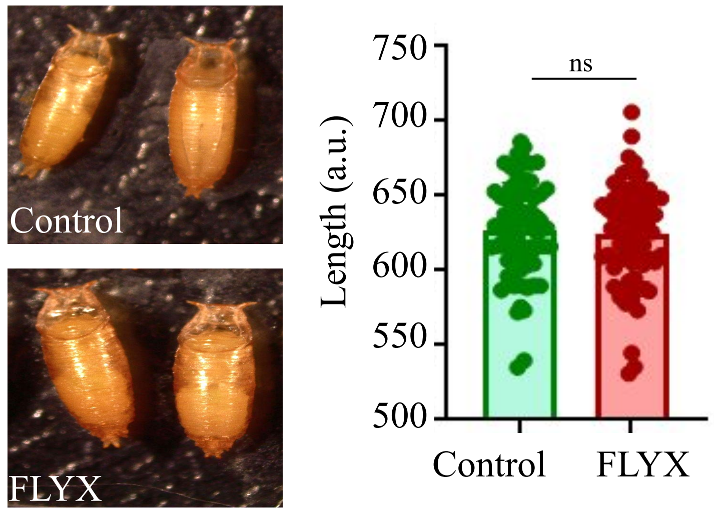


#### Cyto-FLYX does not change fly fecundity

E

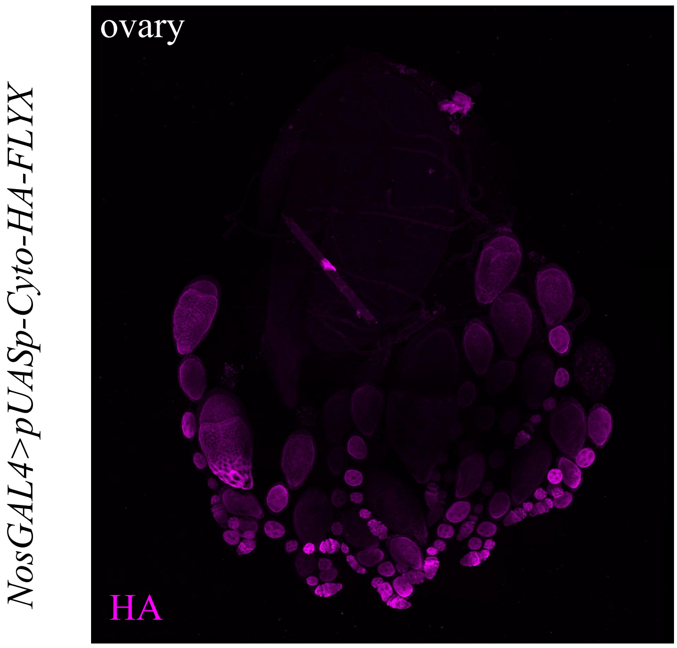


F

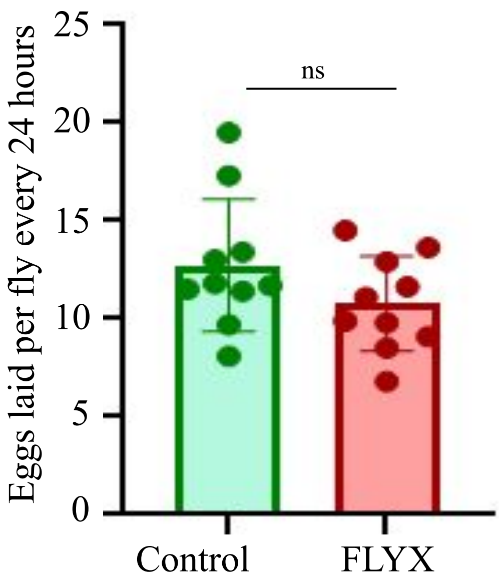

**Supplementary 7**

**A** Cyto-FLYX does not show change in Ecd, InR, and TOR pathway related major transcripts

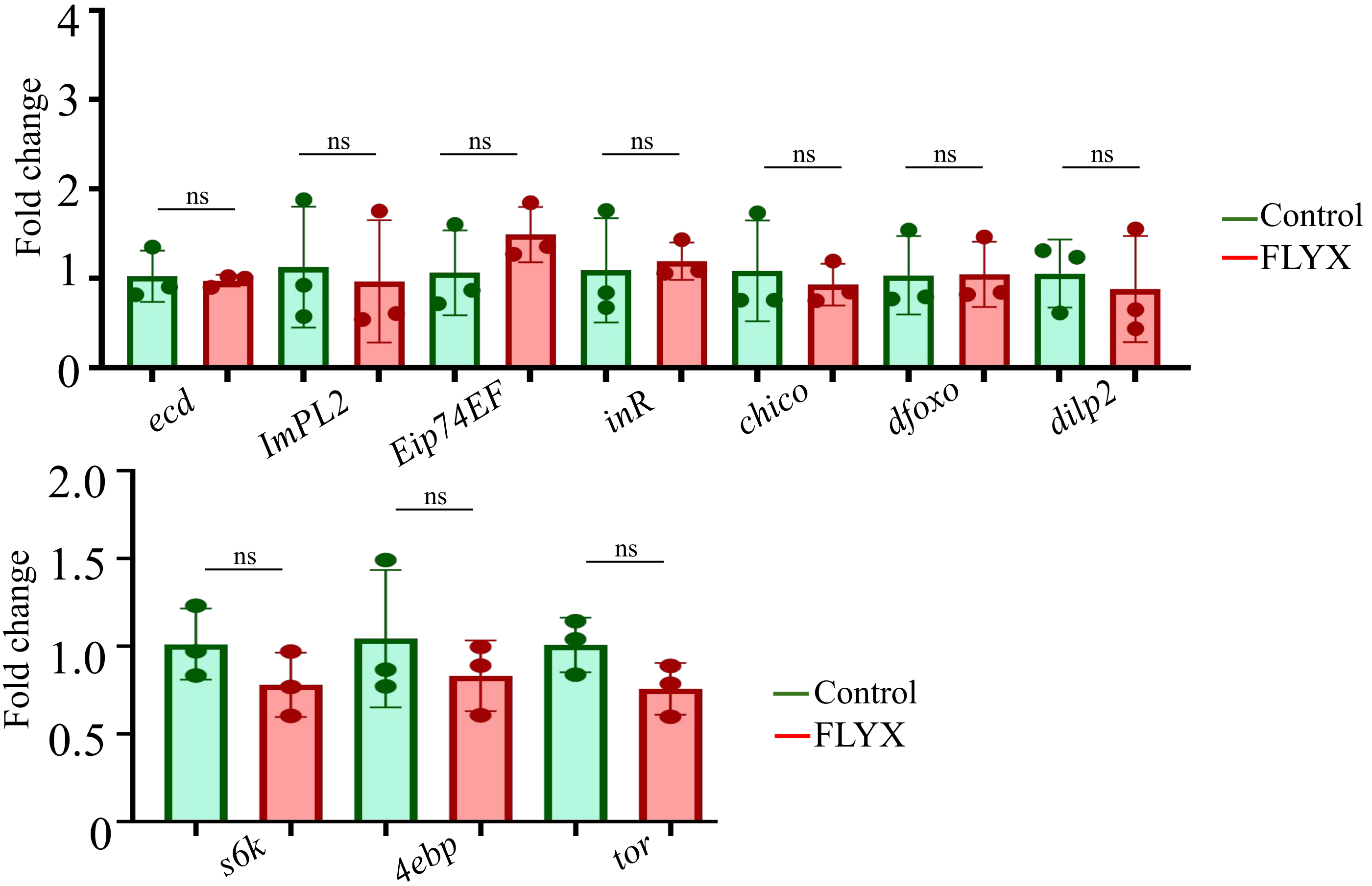


**B** Principal Component Analysis of samples

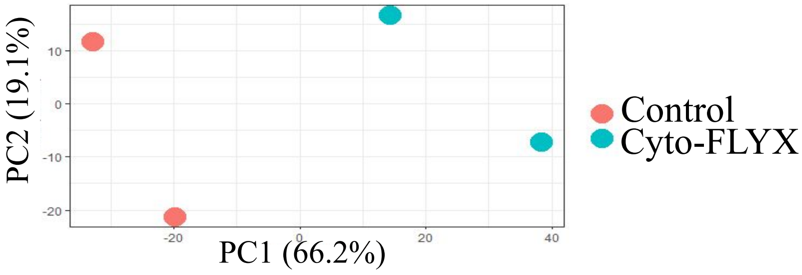

### Supplementary 8

Gene Set Enrichment Analysis in Cyto-FLYX and control larvae

**A** Biological processes

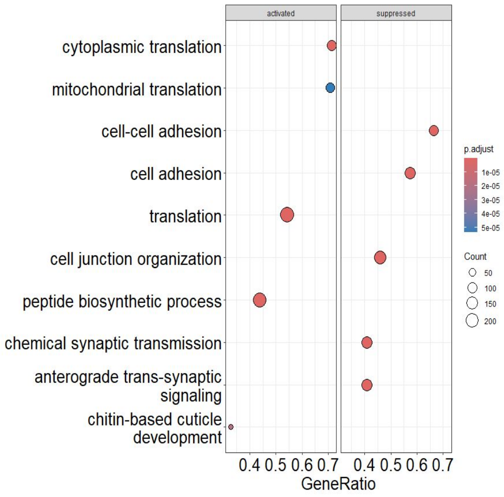


**B** Cellular components

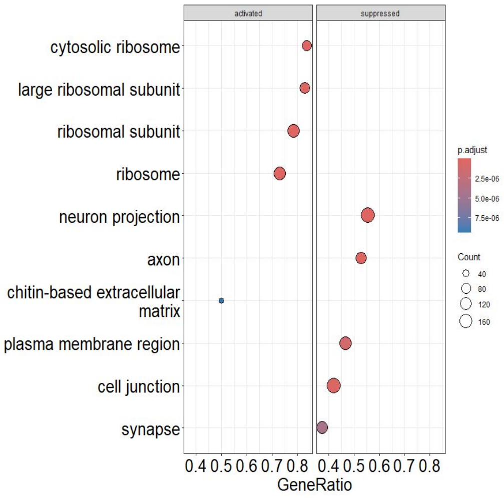


**C** Molecular functions

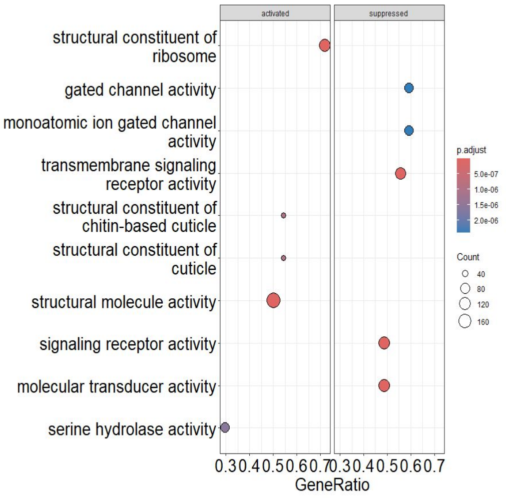


**D** Heatmap of three gene sets enriched in Cyto-FLYX versus control larval RNA samples

Mitochondrial translation

Cytoplasmic translation

Ribosome biogenesis

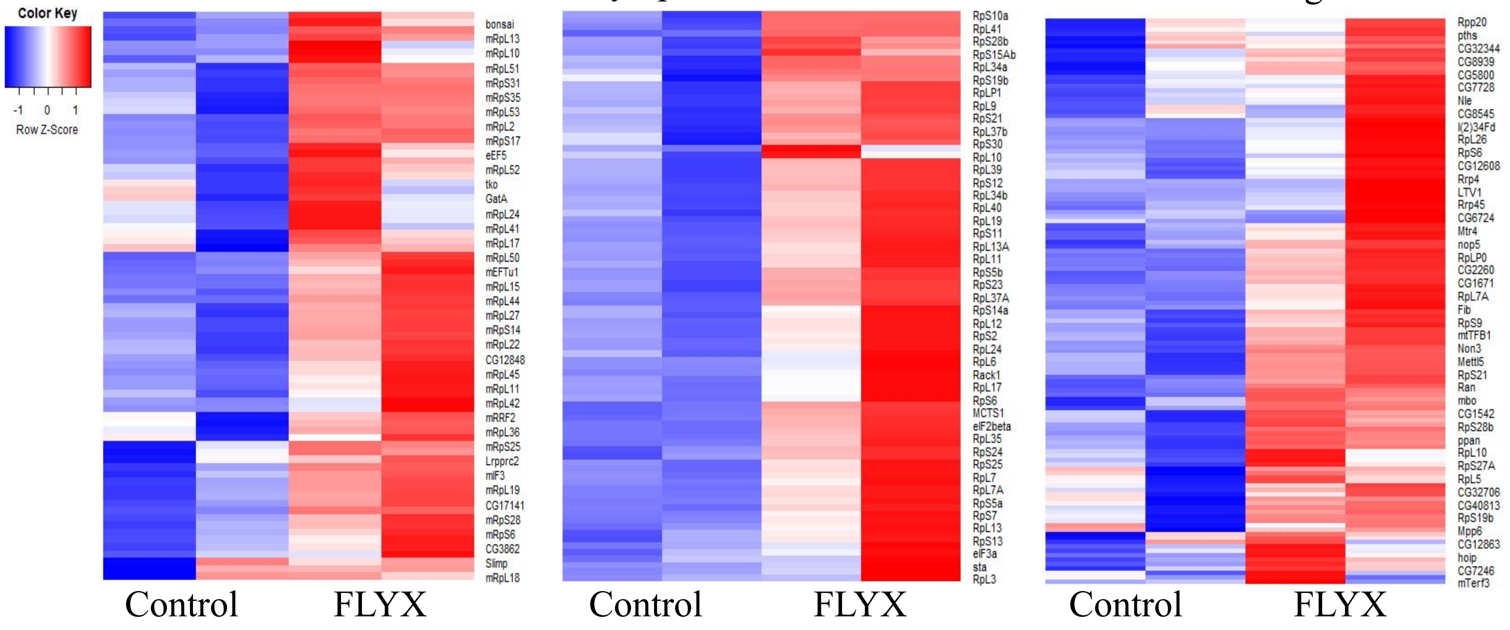
